# PIRD: Pan immune repertoire database

**DOI:** 10.1101/399493

**Authors:** Wei Zhang, Longlong Wang, Ke Liu, Xiaofeng Wei, Kai Yang, Wensi Du, Shiyu Wang, Nannan Guo, Chuanchuan Ma, Lihua Luo, Jinghua Wu, Liya Lin, Fan Yang, Fei Gao, Xie Wang, Tao Li, Ruifang Zhang, Nitin K. Saksena, Huanming Yang, Jian Wang, Lin Fang, Yong Hou, Xun Xu, Xiao Liu

## Abstract

**Motivation:** T and B cell receptors (TCRs and BCRs) play a pivotal role in the adaptive immune system by recognizing an enormous variety of external and internal antigens. Understanding these receptors is critical for exploring the process of immunoreaction and exploiting potential applications in immunotherapy and antibody drug design. Although a large number of samples have had their TCR and BCR repertoires sequenced using high-throughput sequencing in recent years, very few databases have been constructed to store these kinds of data. To resolve this issue, we developed a database.

**Results:** We developed a database, the Pan Immune Repertoire Database (PIRD), located in China National GeneBank (CNGBdb), to collect and store annotated TCR and BCR sequencing data, including from *Homo sapiens* and other species. In addition to data storage, PIRD also provides functions of data visualisation and interactive online analysis. Additionally, a manually curated database of TCRs and BCRs targeting known antigens (TBAdb) was also deposited in PIRD.

**Availability and Implementation:** PIRD can be freely accessed at https://db.cngb.org/pird.

## BACKGROUND

T cells and B cells, as the two pillars of the adaptive immune system, have a crucial role in health and disease. T cell receptors (TCRs) and B cell receptors (BCRs) undergo clonal expansion or antibody affinity maturation to react against antigens. The majority (∼95%) of T cells express TCRs composed of α and β chains (αβ T cells), while the rest of the TCRs are composed of gamma and delta chains (γδ T cells). Similarly, the BCRs expressed on the B cell surface consist of heavy and light chains. During the development of T and B lymphocytes, recombination of variable (V), diversity (D) and joining (J) gene segments (a range of V, D and J gene segments ranked on chromosome), nucleotide deletion at the end of gene segments and nucleotide insertion during the V-D or D-J junctions (V-J junction for α, γ and light chains) cause a highly diverse array of TCR and BCR repertoires (Liu and Wu, 2018). Theoretically, more than 10^18^ unique TCRs and more than 10^13^ unique BCRs have been predicted for human beings (Georgiou, et al., 2014; Venturi, et al., 2008). Therefore, the extremely large diversity of receptors promotes the ability of the immune system to maintain homeostasis in health and disease.

With the advances in high-throughput sequencing platforms, there has been a rapid accumulation of TCR and BCR repertoire sequencing data in the past decade. Specifically, since several influential and seminal studies published in 2009(Boyd, et al., 2009; Robins, et al., 2009; Weinstein, et al., 2009), an increasing number of studies and applications have occurred in subsequent years, which included (i) investigating the immune microenvironment in tumours to assist in therapy by searching for tumour-infiltrating T cells targeting neo-antigens (Reuben, et al., 2017; Riaz, et al., 2017; Wang, et al., 2017; Zhang, et al., 2017; Zheng, et al., 2017); (ii) detecting minimal residual disease and monitoring immune reconstitution after treatment and transplant(Wu, et al., 2012; Wu, et al., 2016); (iii) exploring immune characteristics in autoimmune and infectious diseases and evaluating the effects of vaccines(Gomez-Tourino, et al., 2017; Jackson, et al., 2014; Jiang, et al., 2013; Parameswaran, et al., 2013; Smithey, et al., 2018; Wang, et al., 2018); (iv) identifying disease-associated clones(Dash, et al., 2017; Emerson, et al., 2017; Glanville, et al., 2017; Huang, et al., 2019); and (v) screening the production of monoclonal antibodies targeting specific antigens and human immunodeficiency virus (HIV) by broadly neutralizing antibody(Cheung, et al., 2012; Jardine, et al., 2016).

TCR and BCR sequencing data are valuable resources; thus, a special database to collect and store these sequencing data is highly needed. Recently, the Adaptive Immune Receptor Repertoire (AIRR) Community released a data standard for sharing and developed the platform iReceptor for querying immune repertoire data(Corrie, et al., 2018; Rubelt, et al., 2017); however, iReceptor only provides a primary search function. Here, we have developed a multifunctional database, PIRD, that stores the TCR and BCR sequence information and provides the functions of data statistics, visualisation and interactive online analysis. Additionally, a database of disease- and antigen-associated sequences, TBAdb, was also deposited in PIRD. PIRD provides the opportunity for other investigators to reuse the data more easily and to compare different aspects of the data. The database may aid in the development of disease treatment, drug research and vaccine designs.

## MATERIAL AND METHODS

### Data Sources

To upload the data to PIRD, three datasets are required, which include the project information, sample information and annotated TCR or BCR repertoire sequences. The TCR or BCR repertoire can be captured for a single chain by multiple PCR(Liu, et al., 2016; Robins, et al., 2009), rapid amplification of cDNA ends (5’-RACE)(Warren, et al., 2011), unique molecular identifiers (UMI) (Khan, et al., 2016; Turchaninova, et al., 2016; Vollmers, et al., 2013) and other methods. To obtain paired chains of receptors, such as TCR α and β chains and BCR heavy and light chains, single-cell RNA sequencing technology(Zheng, et al., 2017) and other developed methods(DeKosky, et al., 2013; DeKosky, et al., 2015; Howie, et al., 2015) can be used. These repertoires are then sequenced by high-throughput sequencers, such as Illumina(Zhang, et al., 2017), BGISEQ(Huang, et al., 2017) and 454(Wang, et al., 2015). The sequences can be processed by commonly used tools, such as MiXCR(Bolotin, et al., 2015), MiTCR(Bolotin, et al., 2013), IMonitor(Zhang, et al., 2015), IgBLAST(Ye, et al., 2013), and Change-O(Gupta, et al., 2015), to generate the annotated information. Both gDNA and RNA of lymphocytes derived from peripheral blood and specific tissues can be used to generate the repertoire. Generally, PIRD only collects data from research studies that have been previously published or that are planned to be published in the future. TBAdb, a relatively independent database deposited in PIRD, contains the antigen and disease-associated TCR and BCR sequences that were derived from previously published literature.

### Database Implementation

The implementation of the web application and database was performed in Django, a high-level Python Web framework. The framework of the search engine and data flows of PIRD was performed using Elasticsearch (a distributed data and real-time search engine), which provides association analysis and real-time and seamless retrieval for all data into the database. PIRD depends on the nodes built by MongoDB clusters and Elasticsearch clusters that standardize and integrate data. Additionally, the development of online analysis and the visualisation platform were based on the MongoDB framework.

## RESULTS

### A Brief Introduction and Data Types In PIRD

PIRD mainly includes five basic datasets, including project information, sample information, raw sequencing data, annotated TCR or BCR repertoires, and TBAdb (Fig. 1). First, the project information contains the description of the project, data processing standards, primers used, sample size, and types of receptors, species, individuals, etc. Importantly, one individual may have multiple samples included in the database; this information is listed here. Second, the sample information is made up of two types of data. One type is the data associated with sample collection, processing and phenotype, such as cell origin, receptor type, sequencing platform, gender, age and race. The other type is the overall statistics of TCR or BCR repertoires, which include total sequence number, unique complementarity-determining region 3 (CDR3) number, diversity index (Shannon index), top 10 clones and frequencies, CDR3 length distribution, and V/J/V-J gene usage. Third, raw sequencing data, deposited in the Nucleotide Sequence Archive (CNSA, https://db.cngb.org/cnsa/), which is a newly developed database to store different kinds of raw sequencing data, can be viewed by clicking the weblink provided in PIRD. Fourth, the dataset of annotated TCR or BCR repertoires is the core resource. The annotation is displayed by each unique sequence, including the information of CDRs and framework region (FR) fragments, VDJ assignment, deletion/insertion nucleotides, paired chains, sequence amount, and mapping information. Last, there is a special independent database, TBAdb, deposited in PIRD.

**Figure 1.**
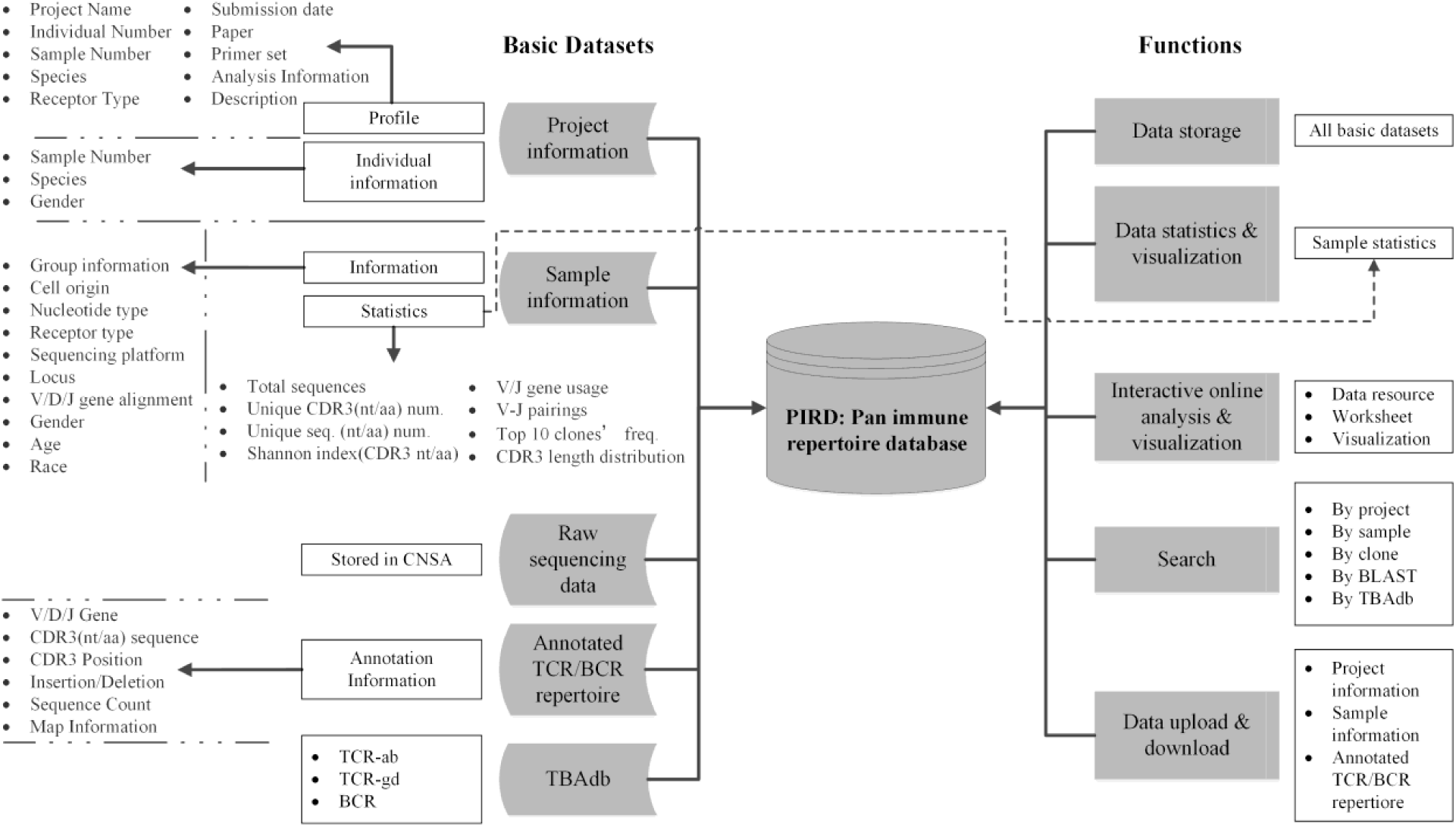
Systematic design of the PIRD database. In this database, there are five basic datasets and five functions. Each dataset or function includes one or multiple parts, and most parts contain several items. Partial items are listed on the left.

### Summary of Data Storage

The data in PIRD are deposited independently by project (Fig. 2B). Thus, the project is the gateway for users to view all data and other information associated with the project (Fig. 2A). Specifically, project information, sample information and raw data can be viewed by clicking ‘ProjectID’. To visit the annotated TCR or BCR repertoire, users can click the ‘SampleID’ shown in the statistics or information pages of ‘sample’. For each sequence, 27 items are used to annotate it by immune repertoire format (IRF-V1.0, https://db.cngb.org/pird/tools/), including VDJ gene assignment, CDR and FR identification, deletion and insertion nucleotides, and mapping information (Fig. 2A). As of July 2019, there are 3657 samples in PIRD, including healthy samples, cancers, autoimmune disease and vaccines, with 877 systemic lupus erythematosus (SLE) samples, 439 healthy samples and 169 leukaemia samples (Fig. 2C). The total number of sequences in PIRD reached 11,395 million (m), and the phenotypes with the top 3 abundant sequences were 2539 m in IgA nephropathy project, 1924 m in minimal residual disease (MRD) project and 1920 m in healthy samples (Fig. 2C). Approximately 76.65% of samples in PIRD are TCR β chain, while others are heavy chain (Fig. 2C). The data in PIRD will be updated over time. In the future, the data will gradually increase as more academic papers are published.

**Figure 2.**
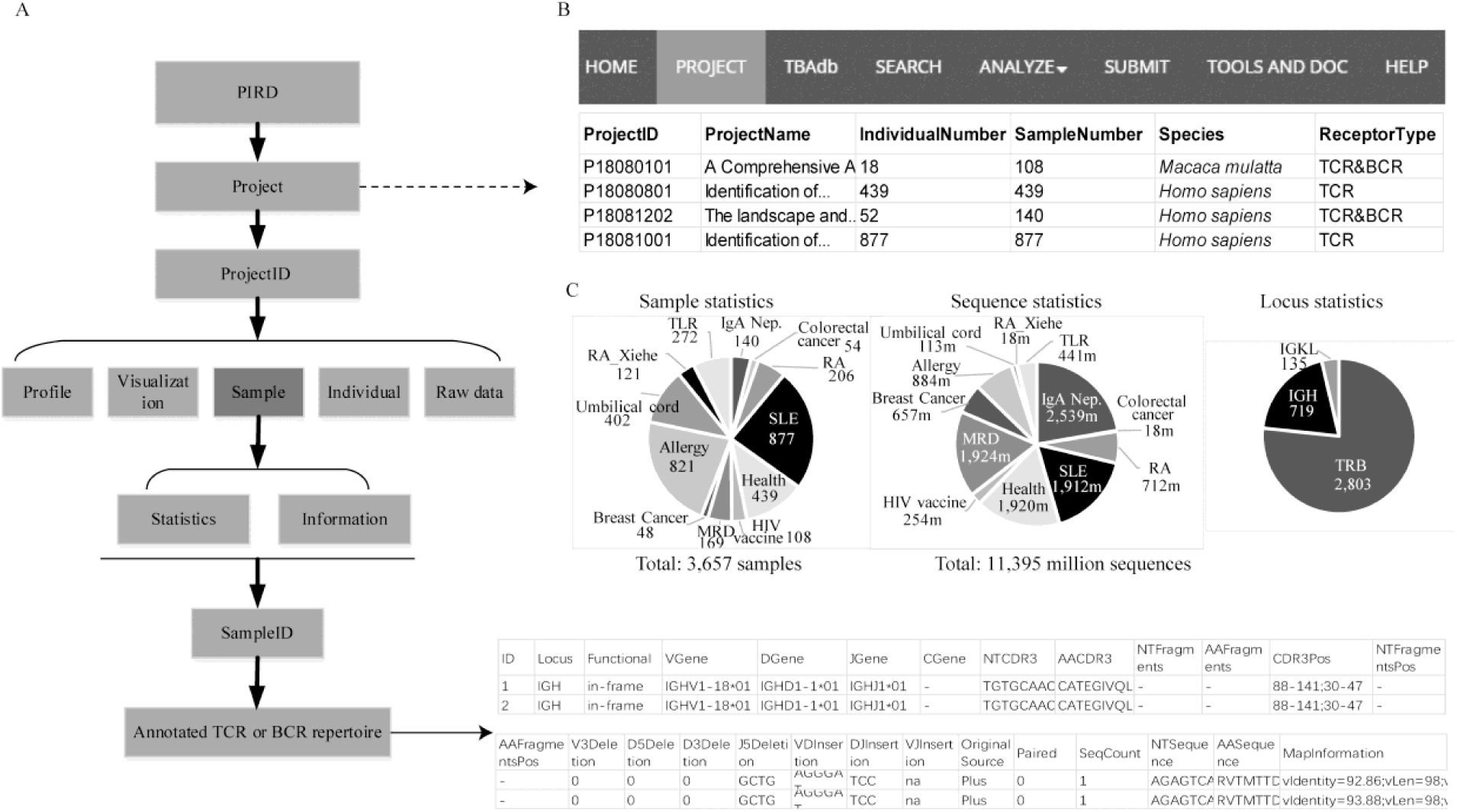
Detailed data deposited in PIRD. (A). Data storage structure for each project. By clicking the links of the PIRD website according to this structure, both sample-associated information and annotated TCR or BCR repertoire can be found. The right bottom corner displays an example of annotated BCR sequences. (B). Project page in PIRD. Partial projects are displayed here. (C). Data statistics in the current database (July 2019). m, million.

### Functions of the Database

PIRD provides data storage, data statistics and visualisation, interactive online analysis and visualisation, search, and uploading and downloading of data (Fig. 1). First, as the fundamental function of PIRD, data storage includes all information of five basic datasets. Second, for each sample, statistics such as CDR3 diversity, CDR3 length distribution, and V/J/V-J pairing usage are calculated from the annotated TCR or BCR repertoires (Fig. 1). To facilitate visualisation by users, PIRD provides corresponding figures with a single click of the button. An example shows one sample’s V-J gene pairing and CDR3 length distribution (Fig. 3A). Additionally, we provide figures for clinical and other data pertinent to the projects, such as the age distribution and CDR3 diversity distribution (Fig. 3B). The third function of the database is the online interactive analysis. We offer a simple and user-friendly platform for users, even those who lack bioinformatics skills, to compare and visualise datasets. For the search function, PIRD provides five types of search modules, including by project, sample, clone, TBAdb and basic local alignment (BLAST) (Fig. 3C). The first three types contain multiple items for users to choose to search for their interested data. The fourth type only searches the data in TRAdb. The last type is a fuzzy search that finds regions of local similarity between sequences using BLAST tool(Altschul, et al., 1990; Ye, et al., 2006) (Fig. 3C).

**Figure 3.**
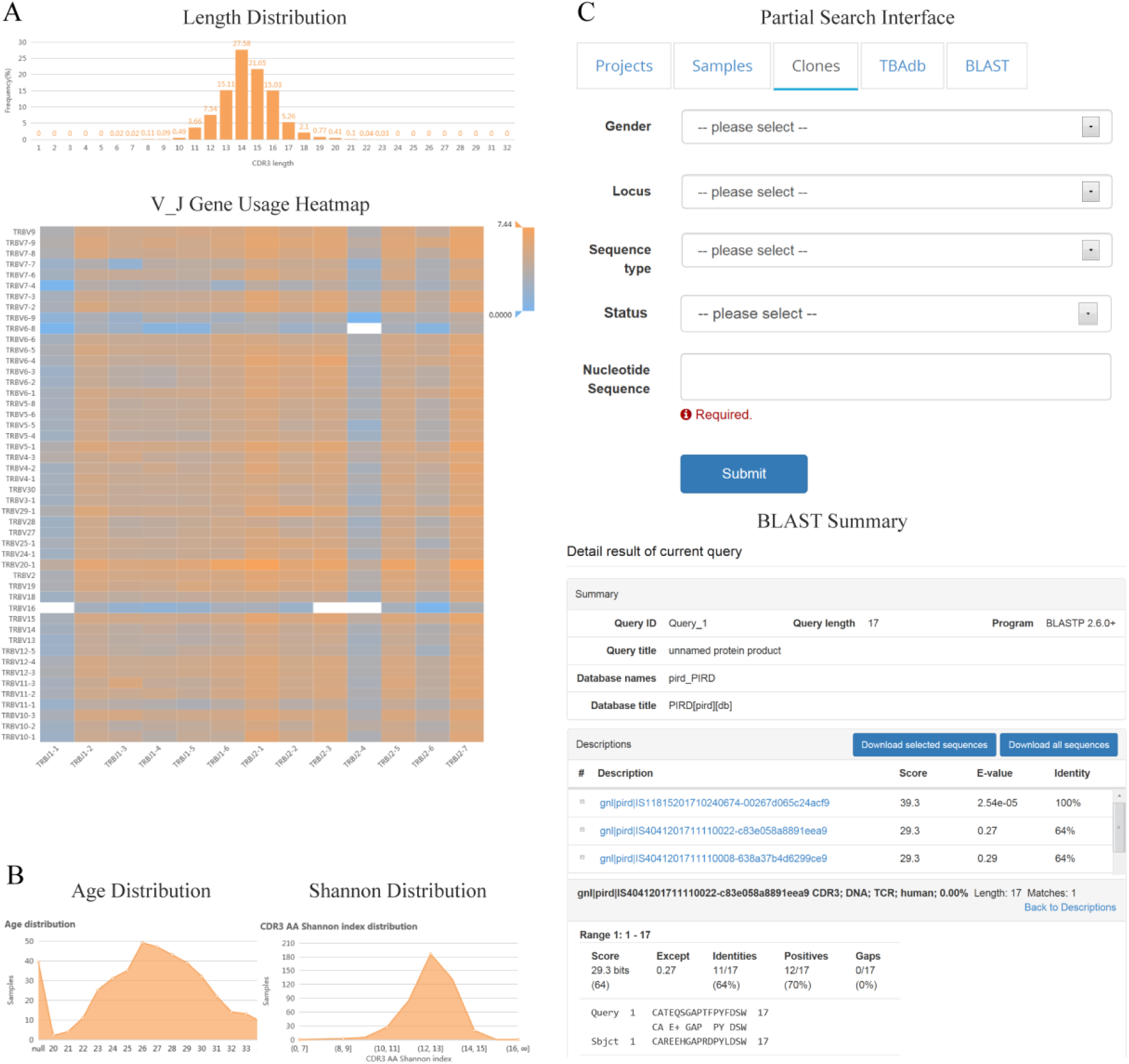
Examples of visualisation and search functions. (A). Visualisation of sample statistics. The length distribution of the CDR3 amino acid sequence (top panel) and the proportion of V-J gene pairings are represented by colour (bottom panel). (B). Visualisation of project information. The left panel shows the age distribution of samples, and the right panel shows the CDR3 diversity distribution in a project. (C). Search interface (top panel) and an example of local alignment search (bottom panel). The data in PIRD can be searched by project, sample, clone and TBAdb, and each provides multiple items to restrict the range of content. The sequences analogous to the query sequences can be found by the local alignment search where the tool BLAST is used.

### Interactive Online Analysis With PIRD

To facilitate direct analysis of data in PIRD, we developed a platform for interactive online analysis. Users can upload their own data or select only specific data in PIRD to create a datasheet. Some analyses, such as comparison of different group features, can be performed, and the corresponding figures, including histogram plot, line plot and box plot, will be generated, which can be performed by dragging the buttons on the left of the website page (Fig. 4A). We used the data of the patient (I190114010005) from a breast cancer-associated project (P19011401) in PIRD as an example and found inconsistent CDR3 length and TCR Vβ gene usage distributions among tumour, adjacent normal tissue and lymph node (Fig. 4B). In addition, we compared the data from multiple projects in PIRD, including those on colon cancer (P18081201), breast cancer (P19011401), healthy people (P18080801) and SLE samples (P18081001). We found that the BCR repertoire diversity of intestinal segment C1 is lower than that of other segments, and the diversity between tumour and adjacent normal tissues is significantly different (Fig. 4C, left panel). Regarding the TCR repertoire, we found that the diversity of SLE is lower than that of the healthy group, and the diversities among different types of samples in breast cancer patients are inconsistent (Fig. 4C, right panel). Overall, the greatest advantage of this analysis platform is that multiple types of diseases or projects deposited in the PIRD can be easily compared.

**Figure 4.**
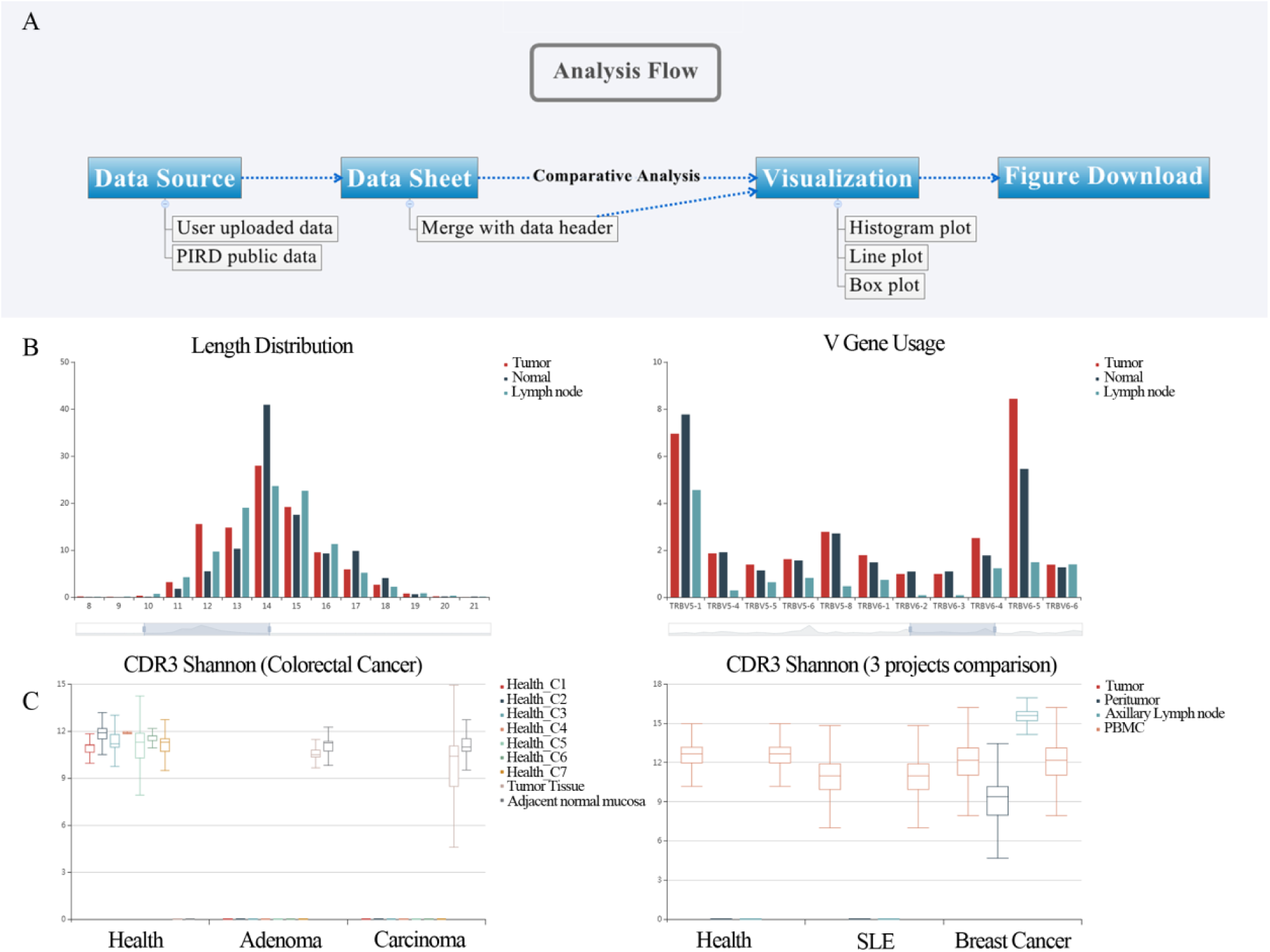
Examples of online interactive analysis and visualisation. (A). Flowchart of the analysis platform. (B). The length distribution of CDR3 amino acid sequences and TCR Vβ gene usage from one patient (I190114010005) with breast cancer (P19011401). There are three samples from the patient, including tumour, adjacent normal tissue and lymph node. All TCR Vβ gene usage can be viewed by sliding the lower box on the website. (C) The diversity of CDR3 amino acid sequences for different groups. The left panel shows the BCR diversities of intestinal samples derived from health, adenoma and carcinoma (P18081201). Each healthy individual contains seven intestinal segments (C1, C2, C3, C4, C5, C6 and C7). The right panel shows the TCR repertoire’s diversities of three projects, including health (P18080801), SLE (P18081001) and breast cancer (P19011401). P19011401 contains samples from tumour, adjacent normal tissue, lymph node and peripheral blood mononuclear cells (PBMC). The diversity is represented by the Shannon index. All data were selected from PIRD. All results of (B) and (C) were obtained using the analysis platform in PIRD.

### Superiorities of PIRD

To date, several repositories have been established to store TCR and BCR repertoire sequences, including ImmuneDB(Rosenfeld, et al., 2017; Rosenfeld, et al., 2018), VDJServer(Christley, et al., 2018) and iReceptor(Corrie, et al., 2018). ImmuneDB and VDJServer mostly focus on online platforms for users to analyse TCR and BCR repertoire sequences. Although users can store the analysed results and share the data in the website of VDJServer, currently, the shared data is very limited. iReceptor is the scientific gateway of the database generated by the AIRR community. Several working groups in the AIRR community, including a common repository group and a minimal standards group, have provided a general standard for sharing the data(Rubelt, et al., 2017). PIRD provides information and data processing pipelines that can be adapted to AIRR standards(Rubelt, et al., 2017). Currently, PIRD provides more functions than iReceptor. First, PIRD provides an interactive online analysis platform, where users can perform some comparison analyses. Second, PIRD offers more statistics and visualisations, such as overall diversities, CDR3 length distribution and V/J gene usage for each sample (ProjectID->Sample->Statistics). Third, for each sequence, the information on deletion and insertion nucleotides, detailed positions of fragments (CDRs and FRs), and paired chains is also included. Fourth, PIRD provides a more powerful search function, including the fuzzy search by BLAST tools. Last, as of July 2019, PIRD has more TCR and BCR sequences than iReceptor (3657 vs. 879 samples, 11,395 m vs. 1,300 m sequences), which is mostly due to a collaboration with the Pan-Immunome Initiative(Xiao Liu, 2018) to release its data before publication. Both PIRD and iReceptor have been trying to provide an application programming interface (API) for users to easily access data from both databases in the near future.

### TBAdb: a manually curated Database of T and B cell receptors targeting known Antigens

Antigen-specific TCRs or BCRs are crucial resources for disease assessment, therapy and vaccine development. To collect previously reported disease and antigen-associated sequences, we read hundreds of published studies and created a knowledge database of TCR and BCR sequences with relevant information. The information collected for each sequence was manually curated and critically checked. The fields include the disease name, antigen, CDR3 sequence, V/J usage, HLA type, experimental method, grade, publication details, etc. Importantly, to evaluate the quality and reliability of antigen or disease specificity, we provide a grade for each sequence according to the identification methods reported in the literature. The criteria for grades are described in the database website (https://db.cngb.org/pird/tools/). TRAdb contains three parts, including TCR αβ, TCR γδ and BCR. It currently includes 52,287 sequences, with 71 diseases. The diseases are divided into five categories, including autoimmunity, cancer, pathogen, allergy and other, like the categories of database McPAS-TCR(Tickotsky, et al., 2017). For the first three categories, each one contains more than 14 diseases (Fig. 5A). In TBAdb, the pathogen category accounts for most of the sequences, which is 89.51% in total (Fig. 5B). We also found that most of the sequences in the pathogen category are more reliable (grade >4, Fig. 5D). This could be because the pathogenic diseases have been widely researched, and the TCR or BCR sequences targeting pathogens are relatively easy to identify. The category with the second largest proportion is autoimmune (Fig. 5B), but more than half of the sequences are shown with a lower grade (Fig. 5D). Although the other category contains 13 diseases, it accounts for only 0.96% of sequences, and most of them are less reliable (Fig. 5D). For TCR αβ sequences, TBAdb contains more data than the VDJdb(Shugay, et al., 2018) and McPAS-TCR(Tickotsky, et al., 2017) databases; however, we found that 9.7% and 5.1% of sequences overlap with the two databases, and 2.3% of sequences are shared among the three databases (Fig. 5C). Therefore, TBAdb is a good supplement for TCR αβ sequences. In addition, TBAdb also includes the data of TCR γδ and BCR.

**Figure 5.**
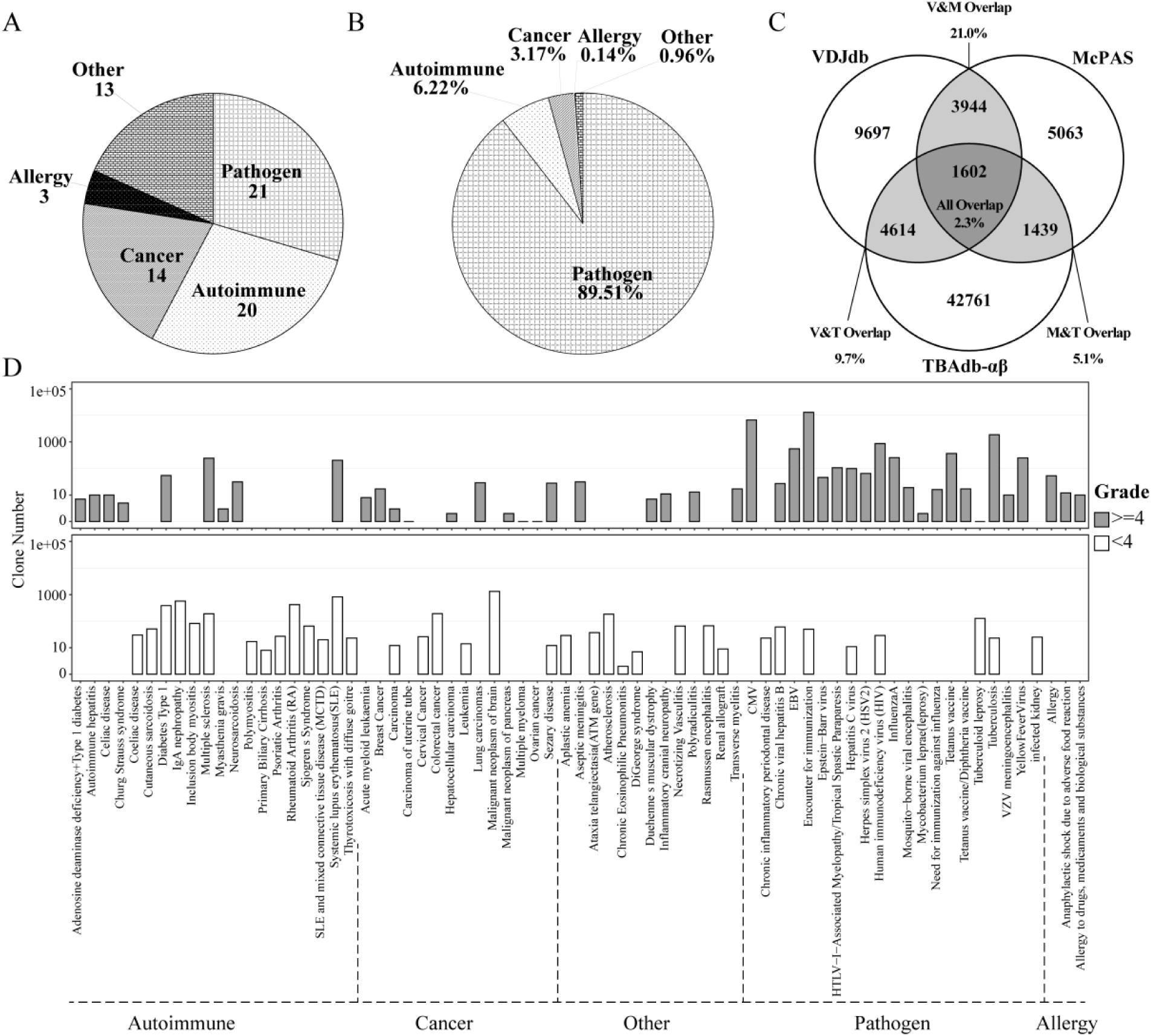
Overall summary of antigen-associated sequences in TBAdb. (A). The number of diseases in five disease categories. (B). The proportion of sequences in each disease category. (C). Comparison of TCR αβ sequences in TBAdb (TBAdb-ab) with VDJdb and McPAS-TCR. V, VDJdb; M, McPAS-TCR; T, TBAdb. (D). The distribution of sequence amount for each disease in TBAdb. The upper panel shows the sequences with grade greater than 4, and the bottom panel shows the sequences with grade less than 4. The grade is a score to evaluate the reliability of sequences associated with the antigen. ‘Other’ signifies diseases that could not be classified into any of the other four disease categories.

## DISCUSSION

Here, we described the PIRD database, a unified repository to store immune repertoire data. The lack of annotated repositories required for immune repertoire sequence storage prompted the creation of PIRD. PIRD is located at CNGBdb (https://db.cngb.org/, Shenzhen, China), which is a newly developed integrative and interactive database. The data in PIRD can be found according to the similarity of sequences from the search page of the CNGBdb. In summary, this multifunctional database provides several benefits to users. First, identifying the disease- or antigen-specific clones is the main purpose of some studies; however, the high diversity of the immune repertoire and very low overlapping rate between individuals(Glanville, et al., 2011) require large-scale samples to evaluate the specificity and sensitivity of the associated clones. Therefore, a wide range of samples in PIRD could provide the function of evaluating clones’ reliability. Second, the bioinformatic methods for immune repertoire data are still in their infancy. Some clustering methods of TCR sequences targeting the same antigen have been recently reported(Dash, et al., 2017; Glanville, et al., 2017), and more machine learning methods will be developed in the future due to the increase in the sequencing data. Thus, the data in PIRD can aid in the development and assessment of new methods by providing training and testing data. Additionally, the sequences in PIRD can also be reused for data mining by the new methods. Third, the online analysis function of the database offers a valuable platform for users to analyse the interested data with a simple interactive interface.

In the future, the online analysis platform in PIRD will be further optimized and developed, including a more friendly interface, more integrative functions and web-based tools. On the other hand, we will continue to collect more sequencing data to store in PIRD. In fact, we have sequenced and processed more than ten thousand samples that will be deposited in PIRD in the near future. Additionally, we will continue to update the TBAdb data in real time. Currently, most sequences in TBAdb are TCR αβ sequences, and since BCR-related studies are increasing, more BCR sequences associated with various antigens and diseases will also be included in future versions.

## CONCLUSIONS

We have developed a database PIRD to collect and store the annotated TCR and BCR sequencing data. It is a multifunctional database, providing data storage and online analysis platform. Other investigators can reuse and compare the data in PIRD.

## ACKNOWLEDGEMENT

This project was funded by the Beijing Genomics Institute and China National GeneBank. The funder provided support in the form of salaries of all authors.

